# Wound induced Ca^2+^ wave propagates through a simple Release and Diffusion mechanism

**DOI:** 10.1101/079053

**Authors:** L. Naomi Handly, Roy Wollman

**Affiliations:** Department of Chemistry and Biochemistry, UCLA, Los Angeles, CA 90095, USA; Institute for Quantitative and Computational Biosciences (QCB), UCLA, Los Angeles, CA 90095, USA; Department of Integrative Biology and Physiology, UCLA, Los Angeles, CA 90095, USA

## Abstract

Damage associated molecular patterns (DAMPs) are critical mediators of information concerning tissue damage from damaged cells to neighboring healthy cells. Adenosine triphosphate (ATP) acts as an effective DAMP when released into extracellular space from damaged cells. Extracellular ATP receptors monitor tissue damage and activate a Ca^2+^ wave in the surrounding healthy cells. How the Ca^2+^ wave propagates through cells following a wound is unclear. Ca^2+^ wave activation can occur extracellularly via external receptors or intracellularly through GAP junctions. Three potential mechanisms to propagate the Ca^2+^ wave are: Source and Sink, Amplifying Wave, and Release and Diffusion. Both Source and Sink and Amplifying Wave regulate ATP levels using hydrolysis or secretion, respectively, while Release and Diffusion relies on dilution. Here we systematically test these hypotheses using a microfluidics assay to mechanically wound an epithelial monolayer in combination with direct manipulation of ATP hydrolysis and release. We show that a Release and Diffusion model sufficiently explains Ca^2+^ wave propagation following an epithelial wound. A Release and Diffusion model combines the benefits of fast activation at length-scales of ~1-5 cell diameters with a self-limiting response to prevent unnecessary inflammatory responses harmful to the organism.

**Abbreviations**
(DAMPs): Damage associated molecular patterns
(ATP): Adenosine triphosphate

## Introduction

The epithelium provides a key protective layer that isolates the internal environment of an organism from outside pathogens. A temporary loss of the epithelial barrier caused by wounds places an organism in a precarious and vulnerable situation (Enyedi and Niethammer, 2015). A timely defense and healing program is vital for organism survival. Wound healing response requires the coordinated action of multiple cell types (Sonnemann and Bement, 2011). Neutrophil cells infiltrate the wounded area to proactively defend against infection. Phagocytosis of pathogens and necrotic cells requires macrophage recruitment. Fibroblasts increase extracellular matrix secretion and provide contractile forces. Finally, epithelial cells proliferate and migrate to close the wound. A plethora of cytokine mediators secreted by the different cell types participating in the wound response regulate the complex wound healing program. However, cytokine secretion, which often requires de novo expression, occurs on an hour timescale (Sonnemann and Bement, 2011; Cordeiro and Jacinto, 2013). The sensitive state of the wounded epithelium requires that wound healing begin as soon as the wound occurs. Therefore, cytokine secretion is too slow to act as the initial signal that activates wound healing programs. A timely wound healing response necessitates a transcriptionally independent program to propagate information regarding the wound to neighboring healthy cells.

Damage Associated Molecular Patterns (DAMPs) is the collective name for a set of chemical ligands that are released from cells upon physical damage (Cordeiro and Jacinto, 2013). DAMPs provide the first indication of damage and are used to activate transcriptionally independent programs. DAMPs quickly propagate information regarding the wound to notify healthy cells surrounding the wound that cellular damage has occurred. Potentially, the gradients formed by DAMPs provide neighboring cells with positional information concerning how far they are from the wound (Sonnemann and Bement, 2011). These damage signals include Ca^2+^ waves, reactive oxygen species, and purinergic molecules such as ATP (Cordeiro and Jacinto, 2013).

Although most known for its key role in cellular metabolism, there has been a growing appreciation for an additional role of ATP as a paracrine signaling molecule (Schwiebert and Zsembery, 2003). Under normal physiological conditions, extracellular ATP levels are typically ~1nM; six orders of magnitude less than cytoplasmic ATP levels of ~1mM (Corriden and Insel, 2010a). This large gradient, actively maintained by cells, makes ATP a powerful damage indicator as any loss of membrane integrity causes an immediate increase in extracellular ATP. Furthermore, previous work has demonstrated ATP as a key initial signaling molecule required for epithelial wound response activation (Feldman, 1991; Caporossi and Manetti, 1992; Pastor and Calonge, 1992).

ATP released from cells following a mechanical wound initiate a Ca^2+^ wave that propagates from the wound in an isotropic pattern. Initially, ATP released from wounded cells binds to P2Y receptors on surrounding healthy cells to increase cytoplasmic Ca^2+^ levels (Figure 1). More precisely, phosphorylation of P2Y by ATP activates PLC to catalyze the degradation of PIP2 to IP3 and DAG (Berridge *et al.*, 2000; Bootman, 2012). IP3 then binds to IP3R on the ER to release Ca^2+^ stores into the cytoplasm. IP3 can pass between cells through connexin channels resulting in intracellular Ca^2+^ activation (Höfer *et al.*, 2001; Warren *et al.*, 2010; Sun *et al.*, 2012; Razzell *et al.*, 2013). While the intracellular pathways that connect extracellular ATP to activate Ca^2+^ signaling have been carefully elucidated (Dupon 2014), the tissue-level pathways responsible for forming the Ca^2+^ wave remain unclear.

**Figure 1.**
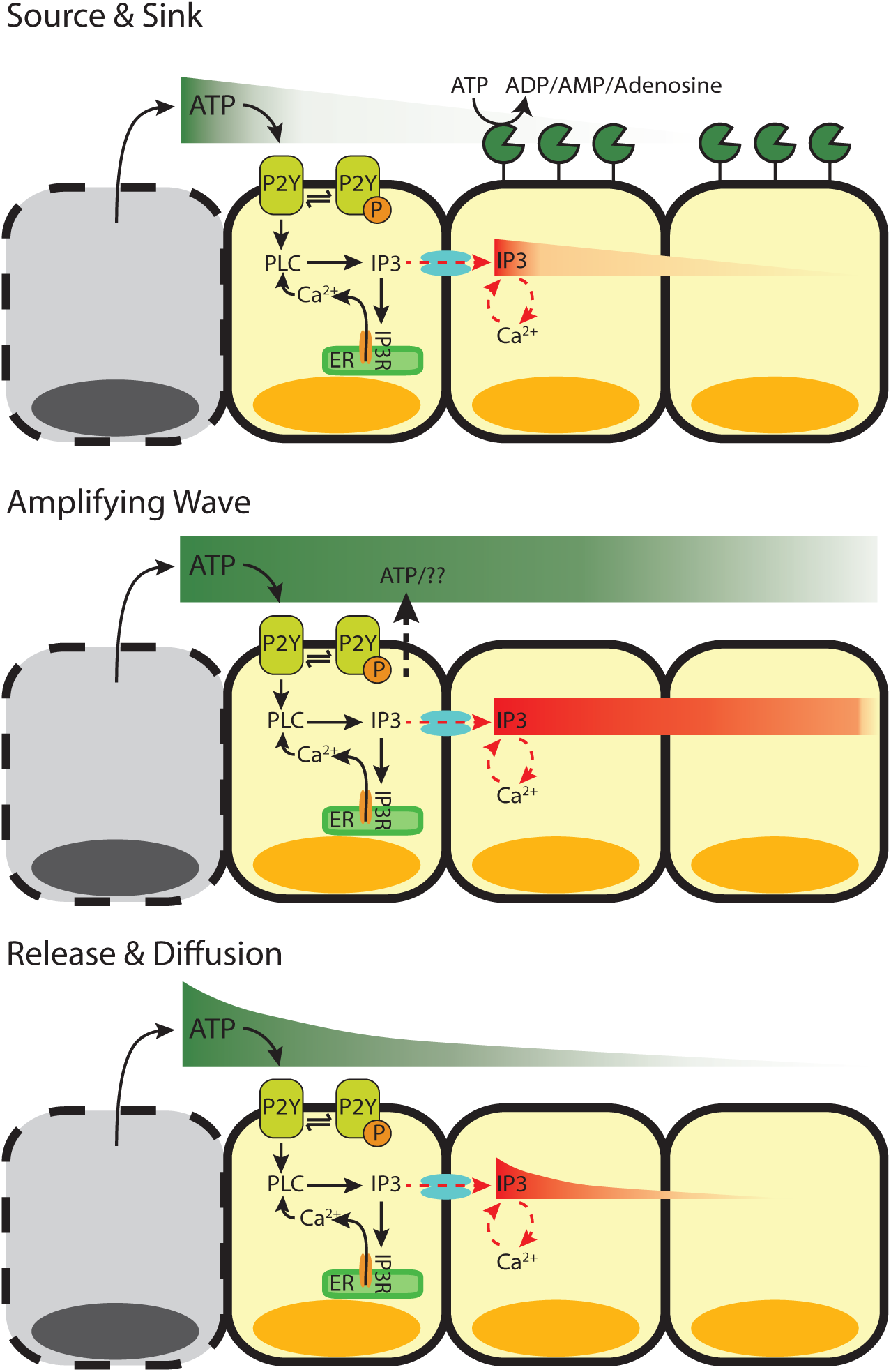
Ca^2+^ wave propagation following wounding. Following wounding, cytosolic Ca^2+^ levels increase in the surrounding healthy cells in a wave-like manner. Ca^2+^ can be activated in cells either extracellularly (ATP binding to extracellular receptors) or intracellularly (IP3 travelling between cells). However, the Ca^2+^ wave can propagate using different mechanisms. 1) Source and Sink. After the initial release of ATP from wounded cells (Source) that activates the cells immediately surrounding the wound, the remaining activating molecules (either ATP or IP3) are hydrolyzed (Sink) as they move away from the wounded area and activate neighboring cells. 2) Amplifying Wave. In this active mechanism, ATP-induced ATP release continues to propagate ATP and IP3 molecules to activate neighboring cells at faraway distances. 3) Release and Diffusion. The release of ATP from wounded cells diffuses away from the wound source to activate neighboring cells with little to no regulation.

Following the passive release of ATP from damaged cells, neighboring cells must be notified of the damage. Several models have been proposed to explain how initial ATP release causes the observed Ca^2+^ wave (Figure 1). In the first model, Source and Sink, ectonucleotidases such as NTPDase2 quickly hydrolyze ATP (Locovei *et al.*, 2006; Ho *et al.*, 2013). In the second model, Amplifying Wave, active propagation mechanisms increase the concentration of extracellular ATP. Specifically, it has been proposed that cells exposed to extracellular ATP respond by actively secreting ATP through Pannexin-1 channels (Locovei *et al.*, 2006; Dubyak, 2009; Corriden and Insel, 2010b; Junger, 2011). In the third model, Release and Diffusion, simple diffusion and dilution of the initial ATP signal controls the Ca^2+^ wave propagation. While molecular studies show support for each model, uncertainty remains concerning which model is primarily responsible to propagate the ATP induced Ca^2+^ wave. Here we use a novel wounding device to mechanically wound an epithelial monolayer (Naomi Handly *et al.*, 2015). Using single-cell wound data combined with genetic and pharmaceutical manipulations, we identify the underlying mechanism responsible for the spread of extracellular ATP in a mechanically wounded epithelial monolayer.

## Results

### Mechanical wounds initiate a Ca^2+^ wave that scales with wound size

Measuring the spatial cellular wound response requires the ability to wound an epithelial monolayer in a convection free environment. We developed a novel microfluidics wounding device to mechanically wound an epithelial monolayer (Figure 2A) (Naomi Handly *et al.*, 2015). Our wounding device has 2 layers: a bottom cell layer and a top air layer. Cells in the cell chamber are mechanically wounded by a pillar in the ceiling of the cell layer when air pressure is increased in the air layer (Figure 2B). Flow is blocked during wounding to prevent any convection within the device to create an isotropic Ca^2+^ wound response (Figure 2C). One advantage of our wounding device over conventional wounding methods such as scratch assays, is the ability to create reproducible wounds (Figure 2D). Reproducible wounding is imperative to measure the spatial response to ensure that each wound elicits a similar response from the surrounding healthy cells.

**Figure 2.**
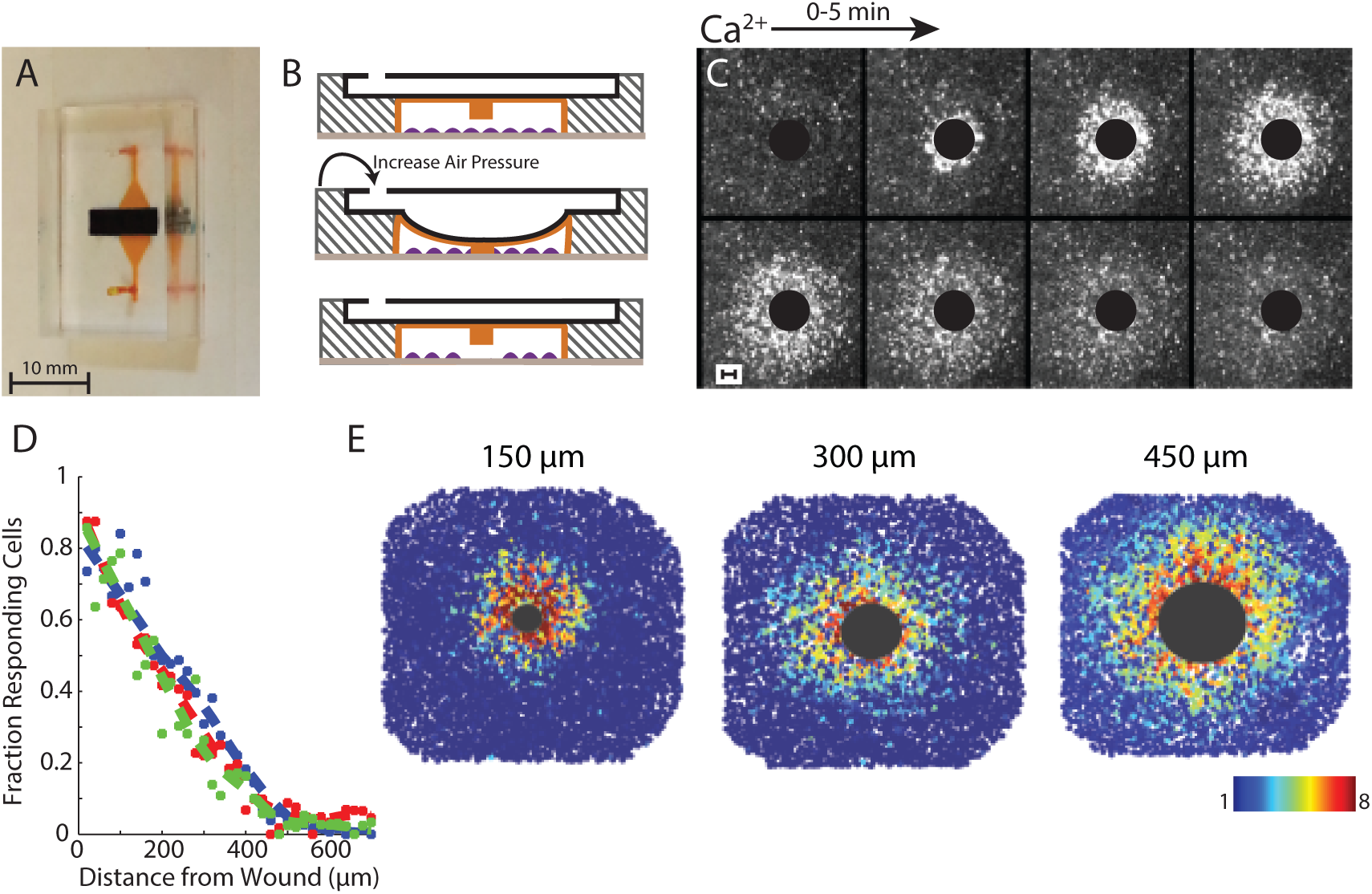
Measuring the Ca^2+^ wave using a microfluidic wounding device. A. Image of dual-layer microfluidics device with the cell chamber (orange) and the air channel (black). B. Cells cultured in the cell chamber (orange) are mechanically wounded when a PDMS pillar in the ceiling of the cell chamber is lowered down when air pressure is increased in the air layer (black). C. Ca^2+^ activation in MCF-10A cells following a 300µm diameter wound. Black circle represents wounded area. Images are over a period of 5 minutes. Ca^2+^ level indicated by Fluo4-AM. D. The fraction of responding cells across the healthy surrounding cells decreases with increasing distance from the wound. E. The Ca^2+^ wave scales with increasing wound size. Large black circle represents wounded area and colored circles represent maximum Ca^2+^ increase per cell. Colorbar represents fold maximum Ca^2+^ increase. Ca^2+^ increase indicated by Fluo4-AM dye.

The isotropic cellular Ca^2+^ response to wounding contains a spatial response that scales with wound size. When measuring the Ca^2+^ response to mechanical wounds, we saw that the fraction of responding cells decreased with increasing distance from the wound (Figure 2D). Additionally, this response scales with wound size (Figure 2E). Spatial information, or knowing where you are in relation to the wound, is imperative during wound healing to ensure that each cell responds appropriately. For Ca^2+^ response we see that cells that are further away from the wound have a smaller and slower response compared to cells that are closer to the wound. Although there are biological processes in place, such as paracrine averaging, to ensure that each cell generates the appropriate response based on its position (Naomi Handly *et al.*, 2015), we wondered how the initial Ca^2+^ gradient forms in an epithelial monolayer following a wound.

### ATP is released in response to mechanical wounding and is required for wound closure

ATP released from wounded epithelial cells is a key damage indicator in the initial wound response (Feldman, 1991; Caporossi and Manetti, 1992; Pastor and Calonge, 1992). We verified whether a monolayer of MCF-10A cells, a breast mammary epithelial line, require ATP release to heal mechanical wounds made by our wounding device. We first established whether initial ATP release is required for a Ca^2+^ response. Indeed, the addition of the ATP scavenger apyrase prevented a gradient of Ca^2+^ activation in cells surrounding the wound (Figure 3A). Next we determined whether the initial ATP released from wounded cells is necessary to the overall wound healing response. Epithelial cells migrate and proliferate in order to close the wound.

**Figure 3.**
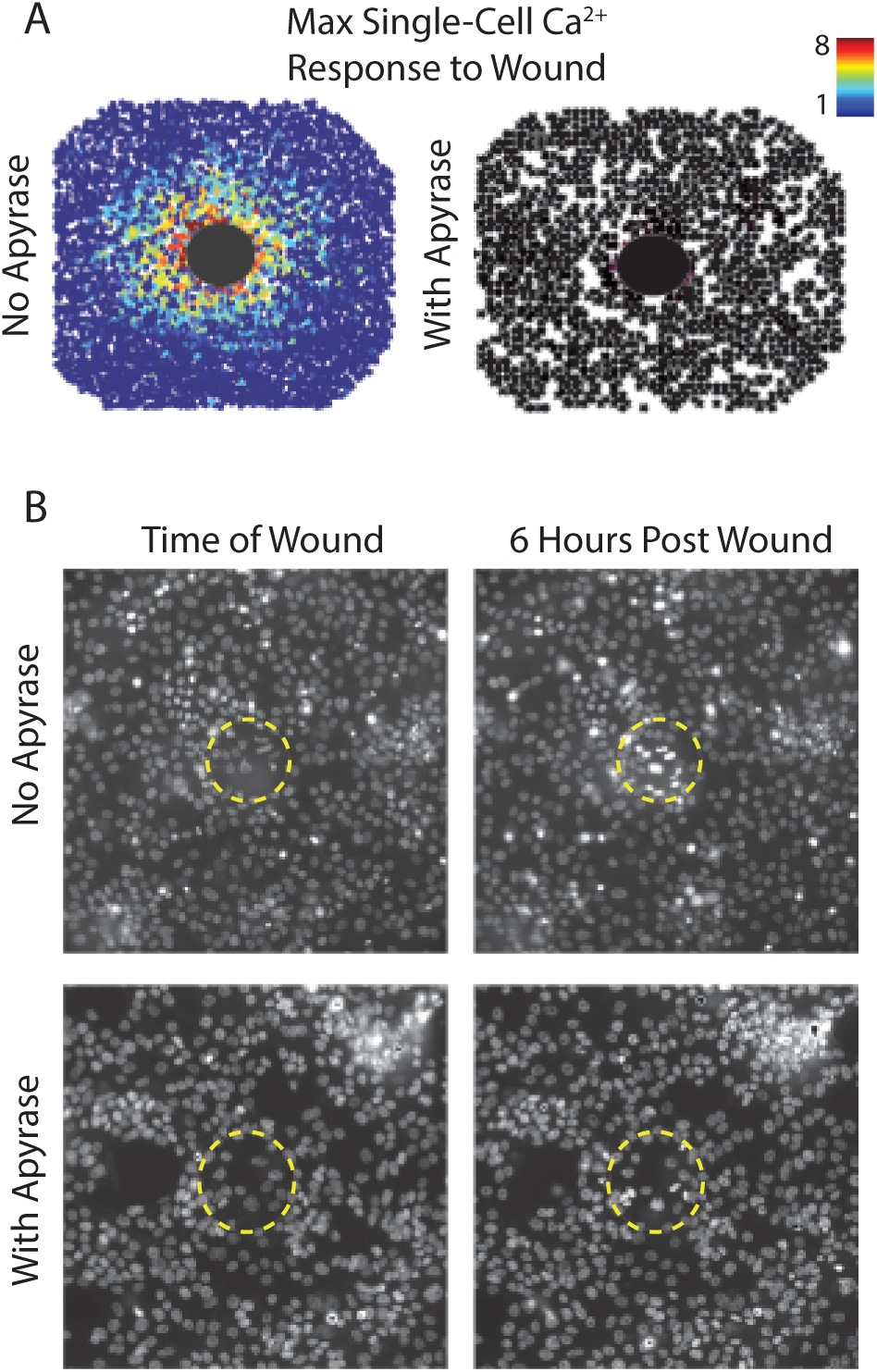
Proper wound closure requires ATP. A. Maximum Ca^2+^ projection indicated by Fluo-4 AM following a 300µm wound in the absence (top) and presence (bottom) of the ATP scavenger apyrase (5U per device). Each circle represents the maximum Ca^2+^ increase per cell and the colorbar represents the fold maximum Ca^2+^ level. Large black circle represents wounded area. Mechanical wounding creates a Ca^2+^ response gradient surrounding the wound. However, apyrase inhibits this response. B. Nuclei with Hoechst following a 150µm wound in the absence (top) and presence (bottom) of the ATP scavenger apyrase. Surrounding healthy cells migrate into the wounded area in the absence of apyrase in 6 hours. No migration is seen following wounding in the presence of apyrase.

Other work has shown that epithelial cells simply sense empty space in order to proliferate and migrate into the wounded area (Block *et al.*, 2004; Klarlund and Block, 2011). However, unlike a scratch assay, wounded cells tend to remain in the wounded area using our wounding device. We see that wounding MCF-10A cells in the presence of apyrase prevents cells from migrating into the wounded area (Figure 3B). Overall we see that the initial release of ATP is required for Ca^2+^ activation as well as epithelial wound closure.

### The Ca^2+^ wave propagates by extracellular activation

The wave of Ca^2+^ activation following wounding can spread using extracellular or intracellular stimulus. Previous data indicates that the Ca^2+^ response following wounding depends on the DAMP ATP (Klepeis *et al.*, 2001, 2004; Yin *et al.*, 2007; Block and Klarlund, 2008; Cordeiro and Jacinto, 2013) (Figure 3). ATP initially released from damaged cells binds to extracellular receptors on neighboring cells causing a cytosolic Ca^2+^ increase within that cell. From here, the propagation of Ca^2+^ activation in neighboring cells can occur via extracellular or intracellular mechanisms. Extracellular stimulation results from ATP propagation that can be augmented by either active release of ATP from cells or degradation by nucleotidases (Locovei *et al.*, 2006; Ho *et al.*, 2013). Intracellular stimulation takes place when IP3 travels between cells via GAP junctions to bind to IP3R to release internal Ca^2+^ stores cells in neighboring cells (Höfer *et al.*, 2001; Warren *et al.*, 2010; Sun *et al.*, 2012; Razzell *et al.*, 2013). In intracellular stimulation, ATP released from wounded cells activate the initial Ca^2+^ response in healthy cells immediately surrounding a wound. GAP junctions then propagate the spread of Ca^2+^ activation to cells farther away from the wound. We first determined whether the Ca^2+^ wave propagates internally or externally.

In order to determine whether the spread of the Ca^2+^ wave is activated through intracellular or extracellular mechanisms, we used our novel wounding device to wound cells in the presence and absence of flow. Based on our previous data (Figure 2) we saw that the Ca^2+^ wave traveled isotropically from the wound in the absence of flow. We wondered whether the response would maintain an isotropic pattern if cells were wounded in the presence of flow. If flow has no influence on the shape of the response, then the Ca^2+^ wave most likely propagates internally where it is not influenced by extracellular flow. When wounding in the presence of flow, however, we saw that the response propagated in the direction of the flow (Figure 4A). This indicates that Ca^2+^ wave propagation is not independent of the extracellular environment.

**Figure 4.**
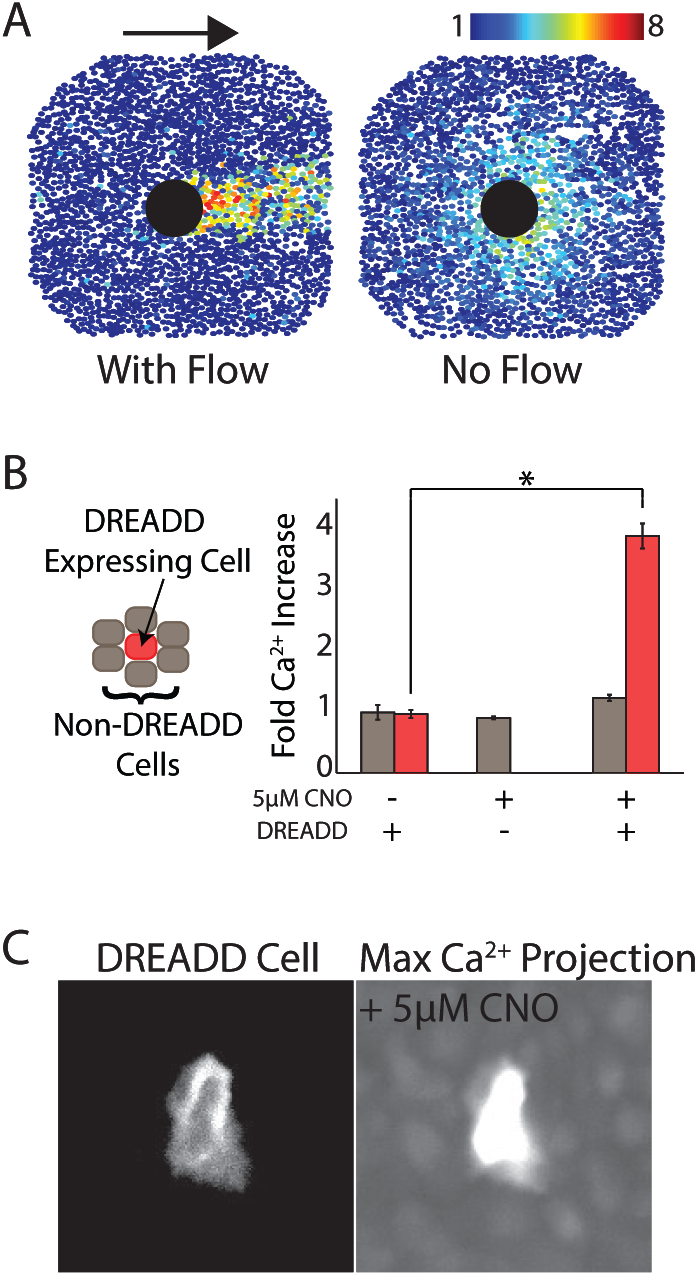
Ca^2+^ wave spread requires extracellular activation. A. When flow is present during the wound, the Ca^2+^ wave goes in the direction of the flow. When flow is blocked, the Ca^2+^ wave propagates in an isotropic manner. Large black circle represents wounded area and colored circles represent maximum Ca^2+^ increase per cell. Arrow indicates direction of flow. Colorbar represents fold maximum Ca^2+^ increase. Ca^2+^ increase indicated by Fluo4-AM dye. B. A co-culture of DREADD expressing (red) and non-expressing (gray) MCF-10A cells show Ca^2+^ activation in DREADD expressing cells only upon the addition of 5µM CNO (error bars indicate SEM, N = 3; *p-value<0.005, t-test). C. Maximum projection of DREADD expressing cell (left) and Fluo-4, AM (Ca^2+^) (right) following the addition of CNO. Ca^2+^ only increases in DREADD expressing cells and not surrounding cells when 5µM CNO is added.

We used a synthetic GPCR DREADD (Designer Receptors Exclusively Activated by Designer Drugs) that is activated by the small molecule clozapine-N-oxide (CNO) to distinguish between extracellular stimulation and artifacts created by flow during Ca^2+^ wave propagation (Armbruster *et al.*, 2007; Dong *et al.*, 2010). We used a DREADD that uses Gq mediated signaling to ensure that Ca^2+^ is activated in cells using the same signaling mechanism as ATP activated Ca^2+^ response. In order to determine whether the Ca^2+^ wave travels internally, we co-cultured DREADD expressing and non-expressing cells such that DREADD expressing cells were surrounded by non-expressing cells (Figure 4B). The addition of CNO activates a Ca^2+^ response in DREADD expressing cells. Whether the Ca^2+^ wave travels internally or externally depends on whether the surrounding non-expressing cells also show an increase in Ca^2+^. A Ca^2+^ response in the surrounding non-expressing cells would indicate an internal propagation mechanism of the Ca^2+^ wave. However, if the wave is dependent on extracellular stimulus, then only the DREADD expressing cells will respond. We found that upon the addition of CNO, only DREADD expressing cells respond. Taken together, these pieces of evidence indicate that the Ca^2+^ wave travels extracellularly.

### Source and Sink: Extracellular ATP degradation does not impact Ca^2+^ activation

In the Source and Sink model, ATP hydrolysis can dominate ATP propagation. Here the lifetime of an ATP molecule determines how long and far it will diffuse from the wound source. Ectonucleotidases present on the plasma membrane metabolize ATP to ADP, AMP, or Adenosine (Idzko *et al.*, 2014). The degradation of ATP released from wounded cells can lead to the formation of the Ca^2+^ wave gradient. In this case, each subsequent cell receives a lower dose of ATP resulting in a lower magnitude of Ca^2+^ activation in a cell. We used pharmaceutical manipulation to determine the role of ATP degradation in creating the Ca^2+^ wave gradient.

We first determined whether ectonucleotidases play a role in ATP mediated Ca^2+^ activation. We used the ectonucleotidase inhibitor ARL67156 at 200µM and added low doses of ATP (0.5 µM). Low concentrations of ATP were used to ensure that cells were not saturated with ATP, making the inhibition of nucleotidases negligible to the overall response. We found that inhibiting ectonucleotidases did not have an impact on the average Ca^2+^ response (Figure 5A). Furthermore, at high levels of ATP (5 µM) we saw no difference between ATP and the nonhydrolyzable analog ATPγS, further supporting that ATP hydrolysis does not play a role in shaping Ca^2+^ activation (Figure 5B)

**Figure 5.**
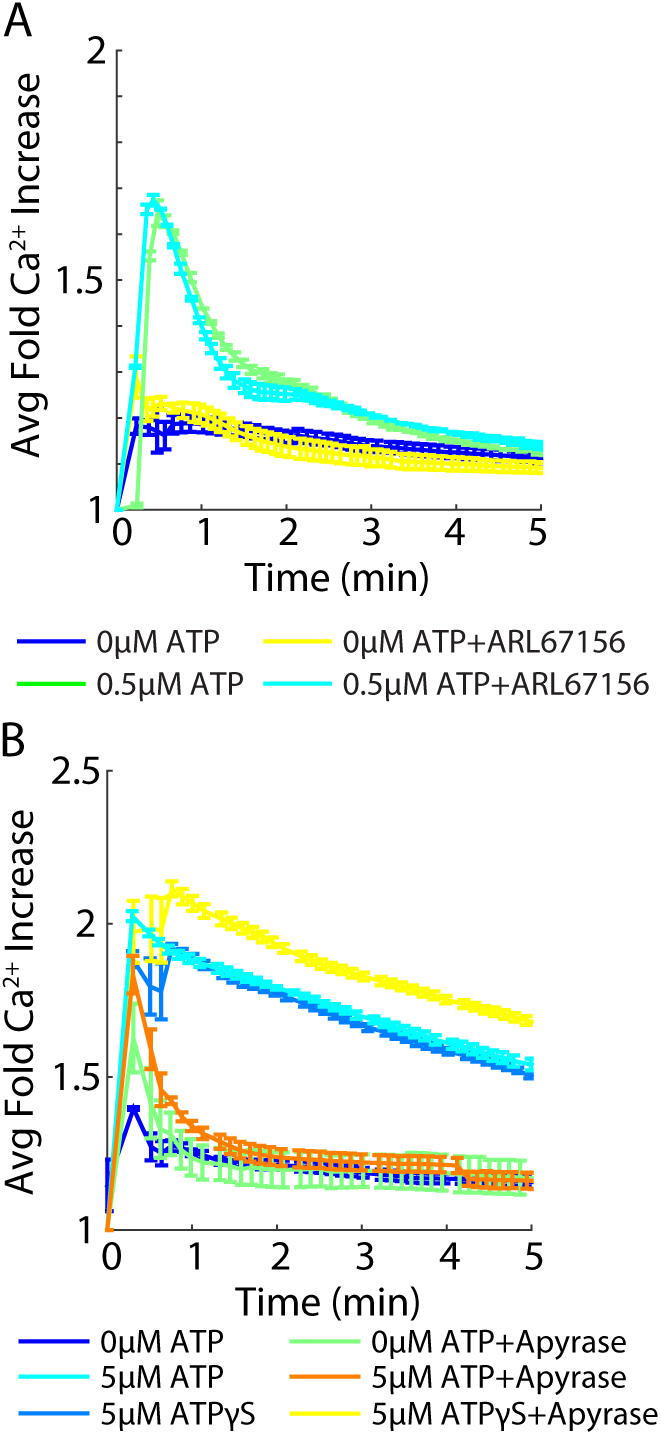
Extracellular ATP degradation does not influence Ca^2+^ activation. Ca^2+^ activation measured by genetically encoded biosensor RGECO. A. Addition of 200µM of the ectonucleotidase inhibitor ARL67156 does not increase Ca^2+^ activation by ATP in MCF-10A cells. (Error bars indicate SEM, n=3). B. Ca^2+^ activation by ATP decreases in the presence of the ATP scavenger apyrase (5U). However, Ca^2+^ activation by the nonhydrolyzable ATP variant ATPγS does not change in the presence of apyrase (Error bars indicate SEM, n=3).

### Amplifying Wave: ATP-induced ATP release does not propagate the Ca^2+^ wave

After determining that the Ca^2+^ wave forms using extracellular stimulus independent of ATP degradation, we determined whether an Amplifying Wave mechanism plays a role in forming the Ca^2+^ wave. That is, does ATP activation initiate the release of ATP to bind extracellular receptors on neighboring cells to propagate the Ca^2+^ response? Prior work has suggested that ATP induces the release of additional ATP from cells (ATP-induced ATP release) (Locovei *et al.*, 2006; Dubyak, 2009; Corriden and Insel, 2010b; Junger, 2011). We first determined whether ATP-induced ATP release is required for ATP mediated Ca^2+^ activation. We perturbed cells with the nonhydrolyzable ATP variant ATPγS in the presence and absence of the ATP scavenger apyrase (Figure 5B). ATPγS activates a Ca^2+^ response in cells similarly to ATP. In the presence of apyrase, any additional release of ATP is hydrolyzed without affecting ATPγS. Apyrase does not deplete Ca^2+^ activation in cells when perturbed with ATPγS as it does with ATP. This indicates that, if an active propagation mechanism exists, cells do not release ATP to activate neighboring cells.

While we saw no evidence for ATP-induced ATP release, it is possible that ATP induces the release of another molecule that propagates the Ca^2+^ wave. We verified the presence of an active release mechanism by measuring the spatial Ca^2+^ activation patterns. Since Ca^2+^ response to wounding is ATP-dependent (Figure 3), we simulated wounding using a photoactivated ATP (NPE-caged ATP) (Figure 6). UV light illumination releases ATP at the site of illumination. We used NPE-caged ATP in combination with a photoactivated fluorescein, CMNB-caged carboxyfluorescein (caged FITC), such that the free ATP and free FITC were released in the same area at the same time. We then compared the spatial activation of Ca^2+^ in the surrounding cells with the diffusion pattern of FITC in a flow-free environment (Figure 6A). Cells were plated in long Ibidi channels of 400µm height to capture the full response length. NPE-caged ATP and caged FITC have similar molecular weights (700.3 g/mol and 826.8 g/mol, respectively). Therefore, the respective molecules will have similar diffusion rates. We reasoned that if an active propagation mechanism exists, the spatial distance of Ca^2+^ activation will extend beyond the diffusion pattern of caged FITC. Furthermore, one port of the channel was sealed prior to uncaging to create a convection-free environment that ensured any response was not due to flow. However, we saw that the Ca^2+^ activation pattern and the caged FITC diffusion pattern are the same in an epithelial monolayer (Figure 6A). This indicates that an active propagation mechanism is not responsible for the Ca^2+^ wave but instead points to a Release and Diffusion mechanism.

**Figure 6.**
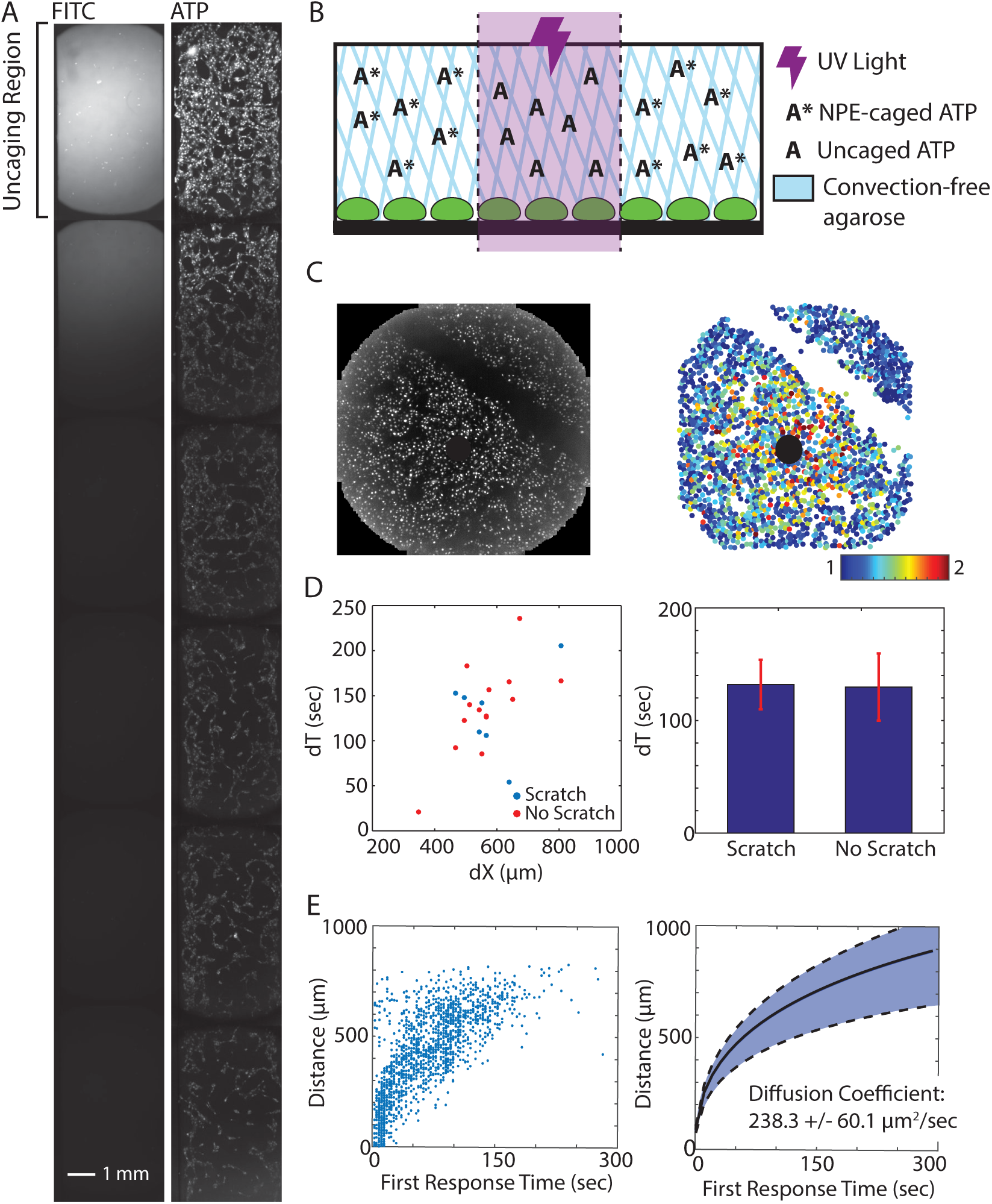
The Ca^2+^ wave propagates by ATP diffusion. A. Photoactivated ATP (NPE-caged ATP) and FITC (CMNB-caged carboxyfluorescein) are uncaged using UV light in the area labeled “uncaging region”. The Ca^2+^ wave, as measured by the genetically encoded biosensor RGECO (right), travels the same distance as uncaged FITC (left) in the absence of any flow. Each side of the channel was taped and allowed to set for 30 minutes to create a flow-free environment. B. Spatially activating cells with ATP in a flow-free environment. An NPE-caged ATP solution is made in 1% agarose and added to cells prior to uncaging. After the agarose has solidified (~10 minutes), a UV light shines on a specific area of the well to release the caged ATP. This spatial increase of ATP concentration mimicsATP released from a wound source. C. The time required for the Ca^2+^ wave to cross empty space (scratched) is measured following photoactivated ATP release (black circle). Right: Raw maximum projection of Ca^2+^ response (measured by RGECO) in MCF-10A cells. Left: Maximum Ca^2+^ projection where each circle represents a cell and the color indicates fold Ca^2+^ increase (colorbar). D. The Ca^2+^ wave takes the same amount of time to traverse a specific distance in the presence (no scratch) or absence (scratch) of cells. E. Propagation of the Ca^2+^ wave according to the first response following ATP uncaging (left) follows a pattern best fit by diffusion. Each point represents the averaged first response time of cells at a given distance following ATP release. The diffusion coefficient that best describes the fitted data is 238.3 +/-60.1 µm^2^/sec (SEM, left, n=3).

### Release and Diffusion: ATP passively creates a Ca^2+^ gradient following wounding

The similar patterns created by FITC diffusion and the Ca^2+^ wave indicate that a Release and Diffusion mechanism may be responsible for Ca^2+^ wave activation. That is, the initial release of ATP from wounded cells diffuses out from the wound to activate neighboring cells. However, other mechanisms may account for these matching patterns. One potential explanation may be that ATP interacts with cells (by binding/releasing to/from extracellular receptors) to form an activation pattern similar to diffusion. In order to determine the presence of an ATP/cellular interaction mechanism, we again used NPE-caged ATP to release ATP from a point source and measured the Ca^2+^ activation patterns over time and space (Figure 6B). In order to maintain a convection-free environment, the NPE-caged ATP solution was made in a 1% agarose solution that solidified on top of cells prior to uncaging. This way, any Ca^2+^ response from ATP would not be due to flow in the well. We first determined whether Ca^2+^ activation rates changed in the presence (no scratch) or absence (scratch) of cells) (Figure 6C). If ATP/cellular interaction takes place, then the first response time of cells after the scratch will differ from the response time of cells at the same distance from the site of uncaging without the scratch. Because scratch wounds also elicit a Ca^2+^ response in the neighboring healthy cells, we made the scratch more than 2 hours before the experiment to allow Ca^2+^ levels to reset to normal levels. In addition, we saw no difference in Ca^2+^ response between scratched and non-scratched wells after uncaging NPE-caged ATP (data not shown). We then measured the time of first Ca^2+^ response for each cell following uncaging. We quantified the difference in first response times for cells before the scratch (closer to the site of uncaging) and after the scratch (farther from the site of uncaging) to find the difference in Ca^2+^ response times (dT). We used this same method for cells at similar distances from the uncaging site but without a scratch and compared the dTs of Ca^2+^ response time for cells with and without a scratch. We found no difference in the dT of Ca^2+^ response time for cells at the same distances from the site of ATP uncaging despite the presence or absence of a scratch (Figure 6D). This indicates that ATP does not interact with cells during the spread of the Ca^2+^ wave. Taken together, the evidence above indicates that there is no active mechanism required to form the Ca^2+^ wave gradient following wounding in an epithelial monolayer of cells. However, we see that the wave forms using extracellular stimulation. Without an activation mechanism, we next tested whether a simple Release and Diffusion model could explain the propagation of the Ca^2+^ wave. Using our NPE-caged ATP method (Figure 6B), we compared the time that each cell first responds to ATP according to its distance from the source with the diffusion coefficient of ATP (Figure 6E). By fitting our Ca^2+^ response data, we calculated a diffusion coefficient of 238.3 (+/-60.1) µm^2^/sec. This value is very close to literature reported ATP diffusion coefficient values of ~300 µm^2^/sec (Warren *et al.*, 2010). Overall, we conclude that a simple ATP Release and Diffusion model is responsible for forming the Ca^2+^ wave. In a Release and Diffusion model, ATP released from wounded cells quickly diffuses from the site of wounding to activate Ca^2+^ response in the surrounding healthy cells in an epithelial monolayer.

## Discussion

Here we investigate the underlying mechanism responsible for propagation of Ca^2+^ activation from a wound. Multiple mechanisms have been proposed for wound-induced Ca^2+^ waves (Klepeis *et al.*, 2001, 2004; Yin *et al.*, 2007; Block and Klarlund, 2008; Razzell *et al.*, 2013). These different mechanisms can be classified under three architypes: Source and Sink, Amplifying Wave, and Release and Diffusion. Evidence from a combination of pharmacological perturbations and quantitative measurements of Ca^2+^ wave propagation contradicted key predictions of the Amplifying Wave and Source and Sink models. These contradictions suggest that the Ca^2+^ wave propagates via a simple Release and Diffusion process. Specifically, ATP molecules released from wounded cells diffuse to activate Ca^2+^ in the surrounding healthy cells. The natural dilution of ATP molecules resulting from diffusion forms the Ca^2+^ gradient across space.

Each of the proposed models has features necessary for effective wound response but also come with key limitations. In order to initiate a proper wound response, cells need to know 1. The location of the wound and 2. The magnitude of the wound. Neutrophils, macrophages, and epithelial cells use signals indicating the location of the wound to find the wound to fight invading pathogens and close the wound. However, this response needs to scale with the magnitude of the wound. A response that is too small results in ineffective healing. A response that is too large can result in excess scarring and even cancer (Darby and Hewitson, 2007; Eming *et al.*, 2007; Schäfer and Werner, 2008; Feng *et al.*, 2010). Additionally, this information must be propagated quickly to begin the wound healing process. By definition the Source and Sink model is self-limiting and ensures that the Ca^2+^ response remains close to the wound. In this model, the response gradient forms when a large amount of initial signal is released (Source) and then hydrolyzed spatially (Sink). This model tightly regulates the ATP propagation distance.

Although this tight regulation is necessary to control the spatial response, it can potentially limit how well the Ca^2+^ response gradient scales with wound size. Contrary to the Source and Sink model, the Amplifying Wave model allows Ca^2+^ activation to spread far from the wound source on a realistic timescale. However, signal amplification may result in information loss concerning the magnitude of the wound. Furthermore, although spreading the signals to far distances may be valuable to recruit immune cells, without self-limiting the wave, the spread of information to distances far from wound can initiate an undesired inflammatory response. The Release and Diffusion model contains features desired for a proper wound response mechanism. A Release and Diffusion model utilizes ATP molecules released from wounded cells as DAMPs to spread to the surrounding healthy cells. It spreads information on a short timescale, scales with wound size, and is self-limiting to prevent undesired activation of faraway cells. Furthermore, an ATP Release and Diffusion model is a simple and straightforward mechanism that requires little additional regulation.

Although the three models we investigated here were chosen as representative mechanistic archetypes, it is possible that the overall Ca^2+^ propagation mechanism uses a combination of models. For example, the Amplifying Wave and Source and Sink models could be combined to a fourth mechanism that balances the contributions of ATP secretion and hydrolysis. However, our data does not support such a model. Our experiments utilizing a cell gap or “scratch” to measure the rate of Ca^2+^ response spread are independent of the molecular pathway underlying an active propagating wave. Additionally, although our results point to a Release and Diffusion model as the core propagation mechanism, it is possible that Amplifying Wave or Source and Sink components act in parallel. Our experimental analysis of Ca^2+^ response propagation did not find any evidence for such parallel mechanisms. If this is the case, any additional components to the core Release and Diffusion model marginally contribute to Ca^2+^ wave propagation.

One key limitation to our result is that our experiments were conducted in an in vitro setting. It is possible that more complex mechanisms occur in vivo. However, the lack of complexity in our model is informative and can direct future research in an in vivo context. It is possible that a 3-dimensional in vivo model, as opposed to the 2-dimensional monolayer in our system, will result in a different response mechanism. Yet, epithelial layers in many organs such as the cornea are very thin and extend to only three cell layers. Therefore, our results provide a good approximation of in vivo geometry. Another important consideration is that the multiple cell types required for wound healing will change the Ca^2+^ wave dynamics and, therefore, the underlying propagation mechanism. Future work quantifying the spatio-temporal dynamics of Ca^2+^ waves in vivo is required to understand how other cell types are involved. It is clear that our model does not capture the full complexity of wound healing in vivo. Yet, the simplicity of a Release and Diffusion model may be beneficial to ensure surrounding cells are alerted quickly. Future work will determine the extent to which a Release and Diffusion model plays a role in the complexity of in vivo wound healing.

## Methods

### Ca^2+^ Measurements in MCF-10A Cells

MCF-10A cells were cultured following established protocols (Debnath *et al.*, 2003). Before plating cells, each surface was first treated with a collagen (Life Technologies), BSA (New England Biolabs), and fibronectin (Sigma-Aldrich) solution in order for cells to completely adhere, according to established methods. In order to maintain a viable environment, cells were imaged at 32°C and 5% CO_2_. All DREADD experiments were conducted in 96-well plates using extracellular hepes buffer (ECB) to reduce background fluorescence (5 mM KCl, 125 mM NaCl, 20 mM Hepes, 1.5 mM MgCl_2_, and 1.5 mM CaCl_2_, pH 7.4). All wound imaging was done in MCF-10A assay media (Debnath *et al.*, 2003).

Single-cell Ca^2+^ levels during wounding and DREADD experiments were measured using Fluo-4, AM (ThermoFisher F14201). Cells were loaded with 8µM Fluo-4, AM, 1X PowerLoad (ThermoFisher P10020), 2.5mM probenecid (P36400), and 20µM Hoechst for 30 minutes at room temperature in ECB. Cells were washed with ECB to remove any remaining extracellular dye. All other Ca^2+^ measurements were conducted using the genetically encoded fluorescent biosensor R-GECO (Zhao *et al.*, 2011; Akerboom *et al.*, 2013).

### Wounding Device Design, Fabrication, and Wounding Assay

Master molds for the microfluidics based wounding device were created using silicon wafers and layer-by-layer photolithography using established methods (Ferry *et al.*, 2011). A separate mold for both the air layer and cell layer were made using negative photoresists and masks. Chips were made by pouring uncured polydimethylsiloxane (PDMS) onto each mold, allowing the PDMS to harden, and bonding the layers together and subsequently to a glass slide. Cells were loaded into the devices through the inlet port using a 20G needle. During wounding the outlet port was plugged using tape and the inlet port held a reservoir of media to prevent evaporation in the chamber. Wounding was accomplished by increasing the air pressure in the top layer of the device until the pillar made contact with the bottom of the device after which the air pressure was released to raise the pillar back up. Cells were loaded in to the wounding device at a density of 15,000,000 cells/mL using a 20G needle. Following trypsinization and resuspension, cells were put on ice to prevent aggregation. Two o-rings were attached to the device surrounding both the inlet and outlet ports for media reservoirs. Each o-ring was attached using a thin film of vacuum grease. Wounding devices were kept in an empty pipet box filled with water to prevent media evaporation. Cells were allowed to adhere for 18-24 hours before wounding.

### Ca^2+^ activation by DREADD

Cells were plated at a density of 2,000,000 cells/100mm plate and allowed to adhere overnight. Cells were transfected with the Gq-coupled DREADD HA-tagged hM3D with an mCherry tag using a 3:1 ratio of FuGene HD (Promega) to DNA and allowed to incubate overnight (Dong *et al.*, 2010). In order to measure the impact of activating a single-cell, non-transfected cells were mixed with DREADD-transfected cells at ratios of 1:0, 1:1, 1:2, 1:5, 1:7, and 0:1 (non-transfected:DREADD) and plated in 96-well plates at a density of 30,000 cells/well. The following day, cells were loaded with 1μM Hoechst dye for nuclear imaging for 30 minutes for cell segmentation purposes. 5μM clozapine-N-oxide (CNO) (Enzo Life Sciences) was added to each well to specifically activate DREADD cells.

### Manipulating extracellular ATP levels

Two methods were used to manipulate extracellular ATP: 1. Inhibition of ectonucleotideases with ARL67156 and 2. ATP hydrolysis with apyrase. Cells were plated in 96-well plates at a density of 30,000 cells/well and allowed to adhere overnight. Cells were incubated with 200µM ARL67156 for 1 hour after which 0.5µM of ATP was added to the well. Extracellular ATP was hydrolyzed by adding 5U of apyrase (Sigma A7646) to the well prior to the addition of 5µM ATP or ATPγS (Tocris 4080). In the wounding device, 5U of apyrase was added to the inlet port just before sealing off the second port to ensure that the apyrase stayed in the cell chamber. Cells were imaged in FluoBrite DMEM media (ThermoFisher A1896701) with the components necessary for MCF-10A assay media added.

### Spatial measurements of the Ca^2+^ wave

MCF-10A cells were plated in coated ibidi µ-Slide VI^0.1^ chips by adding 20µL of a cellular solution with a density of 1x10^6^ cells/mL to each channel of the µ-Slide. Cells were allowed to settle for 1 hour after which each well was filled with MCF-10A assay media (Debnath *et al.*, 2003). Cells were allowed to adhere overnight. In order to measure the distance of Ca^2+^ wave response, we used 1mM NPE-caged ATP (ThermoFisher A1048) and 10mM CMNB-caged fluorescein (ThermoFisher F7103). NPE-caged ATP and CMNB-caged fluorescein were uncaged in specific regions by shining UV light on the region for 20 seconds. Enough NPE-caged ATP solution was added to the channel to partially fill each well. Each well was taped and allowed to sit for >30 minutes to stop flow through the channel.

All other NPE-caged ATP experiments were done in a 96-well plate in an agarose to prevent any convection during uncaging and imaging. NPE-caged ATP solutions were made in a 1% agarose solution and allowed to solidify for 10 minutes at room temperature prior to uncaging. Each well was exposed with UV light for 10 seconds to uncage NPE-caged ATP. To determine whether the Ca^2+^ wave could cross empty space, cells were seeded in a 96-well plate at a density of 30,000 cells/well and allowed to adhere overnight. Cell monolayers were scratched using a 2µL pipet tip to create empty space. Scratches were done >4 hours prior to imaging to give Ca^2+^ levels adequate time to return to basal levels. A 1% agarose solution containing 10µM NPE-caged ATP was added to each well and allowed to solidify for 10 minutes at room temperature. Each well was exposed with UV light for 10 seconds to uncage NPE-caged ATP.

### ATP Diffusion Fitting

We considered the diffusion of a ATP from a single point to its surrounding neighbors to find the diffusion coefficient *D* of the molecule responsible for Ca^2+^ wave propagation following ATP release. We considered a 2D-like geometry where cylindrical cells, each of height *h*_*c*_ and radius *ρ*, grow in a chamber of total *h*_*f*_ height. We simplify the below analysis by approximating the cell monolayer geometry to a series of “cell cylinders”. We also consider the number of molecules released from a cell *N*_*r*_, the number of molecules needed for detection *N*_*d*_, and the total integration time T. The key results of the required integration time are similar for other comparable geometries (data not shown). Under these conditions one could write the analytical solution of the diffusion equations:

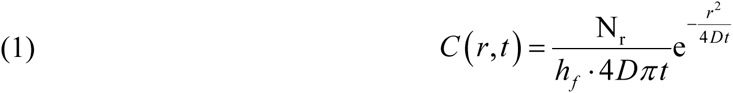

Where *C*(*r*, *t*) is the concentration of ATP for distance *r* and time *t*. For a neighboring cell to respond to this paracrine signal, a critical number of molecules *N*_*d*_ need to reach the volume surrounding the cell. We assume that a cell “senses” a volume comparable to the volume of a cell itself. For a cylindrical cell of area *πρ*^2^ and height *h*_*c*_ the critical concentration required for cellular response will be:

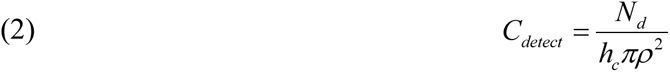

This is simply the required number of molecules divided by the cell volume. Combining equations 1 and 2 we can solve for the distance and time of where the critical concentration will be reached. Solving for distance we get that

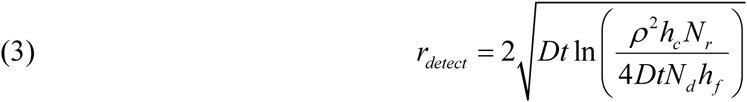

Using the time of first response for each cell with is corresponding distance from the ATP source, we used equation (3) to fit the experimental data to get the ATP diffusion coefficient.

### Imaging and Image Analysis

Imaging was accomplished using a Nikon Plan Apo λ 10X/0.45 objective with a 0.7x demagnifier and Nikon Eclipse Ti microscope with a sCMOS Zyla camera. All imaging was accomplished using custom automated software written using MATLAB and Micro-Manager (Edelstein *et al.*, 2010). Image analysis was accomplished using a custom MATLAB code published previously (Selimkhanov *et al.*, 2014).

## Acknowledgements

The work was supported by GM111404, EY024960 (RW), and a training grant (GM007240) for LNH.

